# Differential glutamatergic and GABAergic responses drive divergent prefrontal cortex neural outcomes to low and high frequency stimulation

**DOI:** 10.1101/2025.03.03.640887

**Authors:** Morteza Salimi, Milad Nazari, Jonathan Mishler, Jyoti Mishra, Dhakshin S. Ramanathan

## Abstract

**Background:** Repetitive brain stimulation is hypothesized to bidirectionally modulate excitability, with low-frequency trains decreasing and high-frequency (>5 Hz) trains increasing activity. Most insights on the neuroplastic effects of repetitive stimulation protocols stem from non-invasive human studies (TMS/EEG) or data from rodent slice physiology. Here, we developed a rodent experimental preparation enabling simultaneous imaging of cellular activity during stimulation in vivo to understand the mechanisms by which brain stimulation modulates excitability of prefrontal cortex.

**Methods:** Repetitive trains of intracortical stimulation were applied to the medial prefrontal cortex using current parameters mapped to human rTMS electric-field estimates. Calcium imaging of glutamatergic (CamKII) and GABAergic (mDLX) neurons was performed before, during, and after stimulation in awake rodents (n=9 females). Protocols included low-frequency (1 Hz, 1000 pulses) and high-frequency (10 Hz, 3000 pulses), with sham stimulation as a control.

**Results:** Glutamatergic neurons were differentially modulated by stimulation frequency, with 10 Hz increasing and 1 Hz decreasing activity. Post-stimulation, 1 Hz suppressed both glutamatergic and GABAergic activity, whereas 10 Hz selectively suppressed GABAergic neurons.

**Conclusions:** These findings provide direct evidence that clinical brain stimulation protocols induce long-term modulation of cortical excitability, with low-frequency stimulation broadly suppressing activity and high-frequency stimulation preferentially inhibiting GABAergic neurons after stimulation.

## Introduction

Repetitive brain stimulation has been used to study synaptic plasticity, particularly long-term potentiation (LTP) and long-term depression (LTD). In 1971, Bliss and Lomo first described LTP, demonstrating that 10-20 Hz or 100 Hz stimulation of the perforant path potentiated synaptic responses in the dentate gyrus ^1^. Follow-up studies, mostly within the hippocampus, helped to refine our understanding of the molecular and cellular aspects that govern LTP^2,3^, including activation of NMDA-receptors, increase in intracellular calcium and AMPA receptor phosphorylation/trans-localization at the synapse ^2,3^. Low frequency stimulation (LFS: 0.5-3 Hz) was later shown to reduce synaptic strength/drive LTD^2–4^ via an NMDA receptor mediated influx of calcium, leading to dephosphorylation and internalization of AMPA receptors^3^. While hippocampal LTP is robust and well-studied, cortical LTP presented distinct challenges: it is difficult to evoke in adults^5,6^, often requires GABA blockade^7,8^, can demand higher-frequency stimulation (>30 Hz)) ^7^ and/or repeated stimulation sessions^9,10^ to induce measurable plasticity.

Repetitive Transcranial magnetic stimulation (rTMS) protocols were inspired by these animal electrical stimulation studies as a method of non-invasively inducing plasticity in humans^11^. Researchers found that high frequency stimulation applied to motor cortex increased the motor evoked potential while low-frequency stimulation decreased the motor evoked potential for about 15-30 minutes ^11–15^. Based on these studies, low-frequency and high-frequency stimulation has been now used in both cognitive neuroscience research^16^ as well as in clinical research and practice for depression^17^, PTSD^18^, headaches^19^, stroke rehabilitation^20^ amongst other conditions. Given its broad clinical use, one might assume that the cellular mechanisms underlying rTMS-induced plasticity are well understood yet surprisingly, they remain unclear. Specifically, how repetitive stimulation alters in vivo neuronal activity and how calcium dynamics during stimulation relate to post-stimulation effects are still poorly defined. Here, we use calcium imaging to examine stimulation-induced plasticity in genetically defined excitatory (CaMKII-expressing) and inhibitory (mDLX-expressing) neurons in the medial prefrontal cortex of rats. We show that clinically observed changes in excitability may be explained by a reduction in GABAergic calcium activity following 10 Hz stimulation and a reduction in glutamatergic activity following 1 Hz stimulation.

## Methods and Materials

### Subjects

This study initially included 14 female Long Evans rats obtained from Charles River Laboratories. However, 5 rats were excluded due to poor signal quality or complications with implant stability precluding recording. The remaining 9 rats included in two groups of CaMKII (5 rats) and mDLX (4 Rats). Female rats were used in this initial study due to initial pilot studies suggesting they easier to record from. Subsequent studies will be performed in males to assess for sex-related differences in effects.

Upon arrival, the rats were approximately one month old, weighing around 150 grams. They were acclimated to the laboratory environment ∼ 4 weeks prior to surgery. During this period, the rats were housed in pairs in standard plastic tubs (10 × 10.75 × 19.5 in, Allentown, NJ, USA). Following surgery, the rats were single housed to facilitate post-operative care and recovery. The housing environment was maintained on a 12-hour light/dark cycle, with lights on at 6 a.m., and all experimental testing was conducted during the light phase. Rats had libitum access to food and water throughout the study to ensure consistent nutrition and hydration. At the start of recording rats were aged between 4 to 8 months and weighed between 250 to 400 grams.

### Electrolens Fabrication

The electrolens assembly (see **Fig 1A)** was constructed by attaching four 50 μm tungsten wires (California Fine Wire) alongside the lens using epoxy. 4 wires were spaced 100 μm apart, starting from beneath the lens edge. The electrodes that demonstrated the most effective stimulation impact on calcium imaging were selected for stimulation delivery. For the ground connection, a 75 μm annealed stainless-steel wire (A-M Systems, Sequim, WA, USA) was used. This wire was soldered to a 5.20 mm × 1.15 mm self-tapping stainless steel bone screw (Fine Science Tools), ensuring secure grounding.

**Figure 1.**
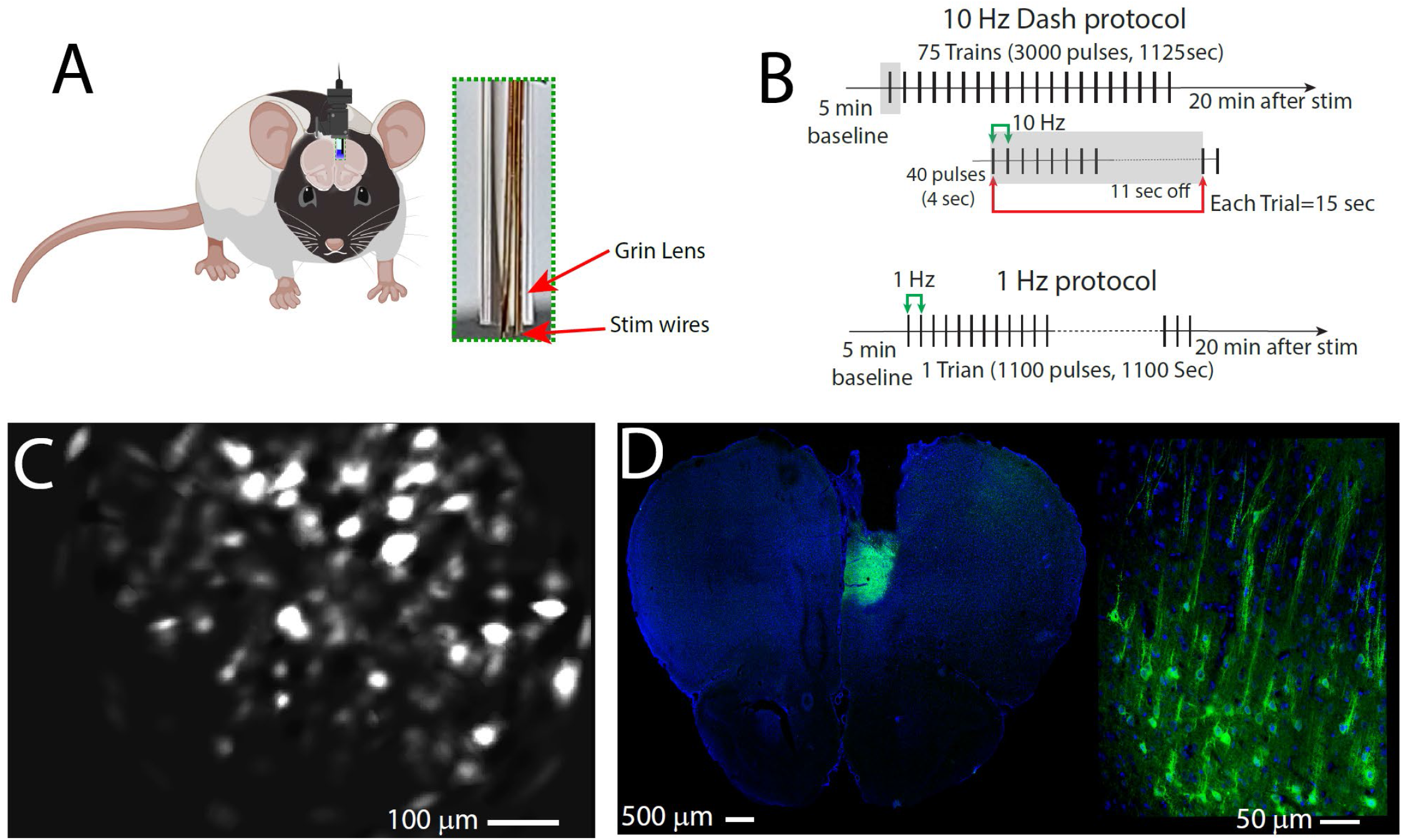
Study Protocol. (A) Schematic representation of the experimental setup showing animals implanted with stimulation electrodes connected to a TDT adaptor for electrical stimulation and an Inscopix miniscope lens for single-cell calcium imaging. (B) Stimulation patterns: In the 10 Hz Dash protocol, 1125 single pulses were delivered across 75 trains, with pulses at 10 Hz for 4 seconds, followed by an 11-second pause before the next train. Each trial consisted of 40 pulses with 15-second inter-trial intervals. In the 1 Hz protocol, animals received 1100 single pulses at 1 Hz without pauses. (C) Screenshot from Inscopix Data Processing software during stimulation. (D) Histological verification of viral expression in the medial prefrontal cortex, visualized in green fluorescence. Left: 10x magnification; Right: 40x magnification.

### Surgical Procedure

Stereotaxic surgery was performed under aseptic conditions to implant recording and stimulation probes. Rats were anesthetized in an induction chamber with 5% isoflurane in 96% room air, delivered using a low-flow anesthetic machine (Kent Scientific, Torrington, CT, USA). Anesthesia was maintained at 1.9–2.5% isoflurane through a nose cone during surgery. A body temperature-controlled heating pad (VWR, Radnor, PA, USA) maintained the rats’ temperature at 37 °C. Rats received a single dose of Atropine (0.05 mg/kg) to reduce respiratory secretions, dexamethasone (0.5 mg/kg) to minimize inflammation, and 1 mL of 0.9% sodium chloride solution for hydration. The surgical site was sterilized with 70% ethanol and iodine solution, and Lidocaine (0.2 cc max) was applied locally. An incision was made to expose the skull, and cranial holes for anchor and ground screws were drilled using a 0.7 mm drill bit (Stoelting). A ground screw was placed posterior to the lambda point, with 3-5 anchor screws secured around the skull periphery using C&B Metabond (Parkell, Inc., Edgewood, NY, USA). Stereotaxic coordinates relative to bregma (AP = +3.72 mm, ML = 0.7 mm) were used to identify the target location in the medial prefrontal cortex (mPFC).

Two viral vectors were used in separate groups of rats: AAV1.Camk2a.GCaMP6m.WPRE.SV40 was injected into the mPFC of five rats to target principal pyramidal neurons. Also, pAAV-mDlx-GCaMP6f6f-Fishell-2(AAV9) was injected into the mPFC of another group of five rats to target inhibitory neurons. The viral vector was delivered using a 33-gauge Hamilton syringe connected to a WPI micro infusion pump at 100 nL/min to a depth of DV = 2.8 mm. The needle was left in place for 10 minutes post-injection to ensure diffusion. A ProView lens (1 mm diameter, 9 mm length) was implanted at DV = 2.4 mm, with a gap between the lens and brain surface sealed using Kwik-Sil (WPI). The lens and electrodes were secured using black dental cement, with connections routed to a TDT ZCA-EIB32 ZIF-Clip® Headstage. Post-surgical recovery was monitored, and analgesics were administered as needed to ensure animal welfare.

This study adhered to NIH guidelines for the care and use of laboratory animals and was approved by the Institutional Animal Care and Use Committee (IACUC) at the San Diego VA Medical Center (Protocol Number A12-021).

### Stimulation Protocol

Electrical stimulation was delivered via a Tucker-Davis Technologies (TDT) 16-channel IZ2H device. Two protocols were applied (**Fig 1B)**. 1)10 Hz Dash Protocol: A total of 3000 single biphasic pulses (cathode leading, 400 µs pulse width, 100µA) were delivered over 75 trains, with train consisting of 10 Hz stimulation for 4 seconds (40 pulses total) followed by an 11-second inter-train interval trains ^21^, a protocol that took 1125 seconds. 2) 1 Hz Protocol: Animals received 1100 biphasic single pulses at 1 Hz with no pauses, a protocol that took 1100 seconds. Sham stimulation consisted of a recording session that was identical to stimulation sessions above but without any stimulation being performed (5-minute baseline, 1125 seconds of no stimulation and an additional 20 minutes of recording after that). Sham stimulation was performed to understand natural dynamics of cellular activity over time.

### Single-Photon Calcium Imaging

Calcium imaging was performed using an nVoke 2.0 miniature microscope (Inscopix) with a 1 mm lens, allowing visualization of neuronal activity in freely moving rats (**Fig 1C)**. Optimal LED power, gain settings, and lens focus were adjusted before each experiment. Imaging was conducted at a frame rate of 20 Hz and processed with Inscopix Data Processing Software (IDPS). Motion artifacts were corrected, and cellular activity was extracted using the CNMF-E algorithm, followed by manual verification to exclude non-cellular signals.

### Processing of Calcium Signals

Data analysis was performed using MATLAB v2023a, ensuring a robust framework for processing and interpreting neuronal activity. Cell classification was based on the Z-score normalization of cellular activity, calculated across the entire recording session. The baseline activity was defined as the average cellular activity recorded during the 5-minute period preceding the onset of stimulation (or sham).

To classify individual cells, their 1 sec average activity during the stimulation (or sham) period was compared to their baseline activity. Cells were categorized into three groups excited, inhibited, or non-responsive based on their Z-score activity during stimulation relative to baseline. Classification was determined using the Wilcoxon rank-sum test, a non-parametric statistical method implemented in MATLAB. A significance threshold of p < 0.01 was applied, ensuring stringent identification of cells with significant changes in activity.

For examining the relationship between neuronal activity during stimulation (or sham) and post-stimulation (or post-sham) periods, a linear regression model was employed. This model was designed to evaluate the consistency and stability of cellular responses over time by analyzing the correlation between Z-score values in these two phases. The regression coefficients and their associated p-values were computed to quantify the strength and statistical significance of the relationships. This analysis provided detailed insights into how neuronal activity patterns evolved following stimulation, highlighting differences in response stability across various stimulation protocols.

### Histological Verification

To confirm viral expression, brain tissue was processed post-experiment (**Fig 1D)**. Rats were perfused with 0.9% saline followed by 4% paraformaldehyde (PFA). Brains were post-fixed in PFA, cryoprotected in 30% sucrose, and sectioned at 40 μm thickness using a cryostat (Leica CM1950). Sections were counterstained with DAPI and analyzed under a confocal microscope (Zeiss LSM 880). Viral expression was verified by green fluorescence signals, matched to anatomical coordinates from the Paxinos and Watson rat brain atlas to ensure accurate targeting of the mPFC.

### Statistical Analyses

All statistical analyses were conducted using SPSS, while data visualization and plotting were performed in GraphPad Prism. Neuronal responses were classified using Wilcoxon rank-sum tests (p < 0.01) to compare activity against baseline. Differences in mean Z-score activity across stimulation conditions were analyzed using one-way ANOVA, followed by Sidak’s post hoc correction for multiple comparisons. Two-way ANOVA was used to assess interactions between time and stimulation conditions, with post hoc tests applied where appropriate. Chi-square tests were conducted on binary-classified data to compare the proportions of excited, inhibited, and non-responsive cells, with post hoc comparisons for significant differences. Linear regression analyses were performed to examine the relationship between neuronal activity during and after stimulation, with R-squared values and Pearson correlation coefficients reported. To control multiple comparisons, family-wise error rate corrections were applied where necessary. All statistical tests were two-tailed, with significance set at p < 0.05 unless otherwise specified. Data are presented as mean ± SEM.

## Results

We found a significant effect of stimulation in the excited cell population (one-way ANOVA, F_(3,1520)_=12.18, *p*<*0*.*0001*; **Figure 2D**). Post hoc analyses showed that 10 Hz stimulation resulted in a significant increase in the mean Z-score activity of excited cells compared to both sham (*p<0*.*0001*) and 1 Hz stimulation (*p<0*.*0001*) while 1 Hz stimulation was not significantly different than sham (*p>0*.*9*). We performed the same analysis on inhibited cells and did not find a significant effect of stimulation (F_(3,1907)_=2.39, *p*=0.066) (**Figure 2E**). Thus, our results suggested that the population effects of 10Hz stimulation could be explained by a preferential increase in calcium activity for cells that were likely modulated in that direction, and did not significantly affect activity in cells that were being suppressed during that time period. More puzzling, however, was that the effects of 1Hz stimulation were not significant for either excited or inhibited populations during stimulation. Thus, we explored an alternate hypothesis that stimulation changes the distribution of responses (i.e. that cells that were likely to be activated in a sham stimulation were suppressed or vice versa). Chi-square analyses (**Figure 2F**) indeed revealed significant differences in the proportion of excited cells (χ ^2^ (3) = 42.6, p < 0.0001) and inhibited cells (χ^2^(3) = 31.09, p < 0.0001). 10 Hz stimulation was not different from sham in the proportion of cells that were excited or inhibited. However, 1Hz stimulation induced a significantly greater proportion of cells that were inhibited and a reduced proportion of excited cells relative to both sham and 10Hz protocols (See Table S1 for significance). Thus, for 1Hz stimulation we found the population level reduction in calcium activity could be explained by the fact that cells that would normally have increased their calcium activity ended up becoming suppressed with stimulation. Overall, these findings demonstrate that 10Hz stimulation can increase calcium influx into glutamatergic cells, thus resulting in a population level activation of the stimulated cortex, while 1Hz stimulation suppresses calcium activity in cells that would otherwise have shown increased calcium levels over time, thus resulted in a net suppression of excitability in the stimulated cortex.

**Figure 2.**
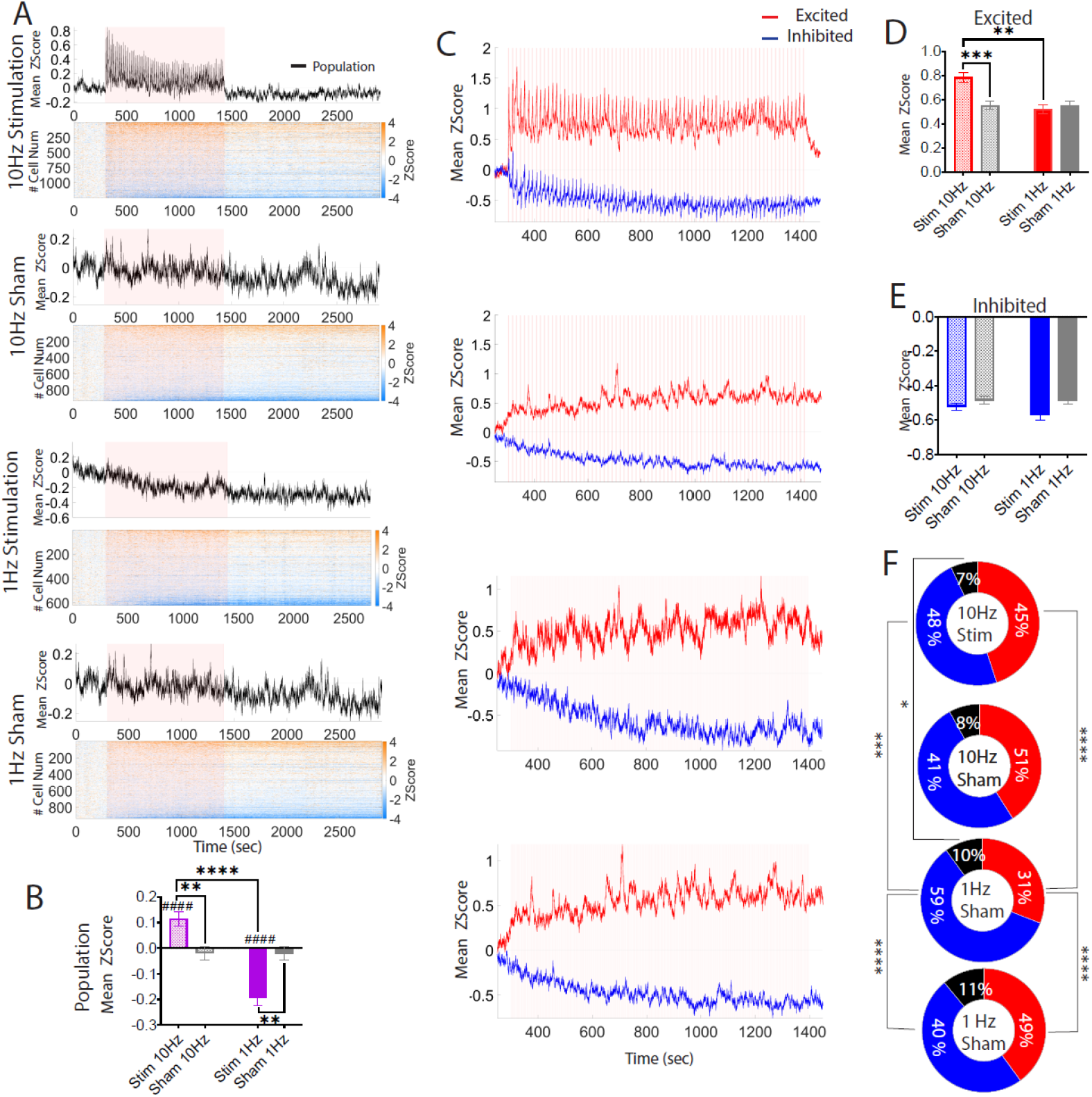
CaMKII Neuronal Responses During Stimulation. **(A)** Average Z-score of overall cellular activity throughout the entire recording session for different stimulation protocols and sham conditions. Z-scores were calculated based on a 300-second baseline period prior to stimulation. **(B)** Mean Z-score activity for sorted cell populations during stimulation. Purple represents the activity of the entire population (including both excited and inhibited cells), red denotes excited cells, and blue denotes inhibited cells. Cells were classified based on significant activity changes compared to baseline (p < 0.01). **(C)** Mean Z-score response of the entire cell population during the stimulation period. Stim 10 Hz (n = 1156), sham 10 Hz (n = 860), stim 1 Hz (n = 562), and sham 1 Hz (n = 857) were compared. One-way ANOVA indicated a significant increase in stim 10 Hz compared to sham 10 Hz (*p = 0*.*0026*) compared to stim 1 Hz (p < 0.0001). Stim 1 Hz also indicated a significant decreased compared to sham (*p=0*.*0012*). One-sample t-tests comparing responses to zero showed a significant increase for stim 10 Hz (p < 0.0001) and a significant decrease for stim 1 Hz (p < 0.0001). **(D)** Mean Z-score response of excited cells during stimulation. Stim 10 Hz (n = 562), sham 10 Hz (n = 365), stim 1 Hz (n = 194), and sham 1 Hz (n = 383) were analyzed. One-way ANOVA showed a significant increase in stim 10 Hz compared to sham 10 Hz (p<0.0001) and stim 1 Hz (p<0.0001). **(E)** Mean Z-score response of inhibited cells during stimulation. Stim 10 Hz (n = 594), sham 10 Hz (n = 475), stim 1 Hz (n = 368), and sham 1 Hz (n = 474) showed no significant differences according to rANOVA. **(F)** Proportions of cell categories (red, excited; blue, inhibited; and black, non-responsive) across different stimulation and sham protocols. Chi-square analyses revealed significant differences in the proportions of excited cells (χ2(3) = 42.6, p < 0.0001), inhibited cells (χ^2^(3) = 31.09, p < 0.0001), and non-responsive cells (χ ^2^ (3) = 9.3, p = 0.025). All data are presented as mean ± SEM. Statistical analyses were performed using rANOVA followed by Sidak’s correction for multiple comparisons. Significance levels: *p < 0.05, **p < 0.01, ***p < 0.001, ****p < 0.0001. ^####^ p < 0.0001 for one-sample t-tests comparing responses to zero.

### Post-Stimulation Effects on Excitability

The canonical model of repetitive brain stimulation suggests that high-frequency stimulation induces long-term potentiation (LTP), increasing cortical excitability, while low-frequency stimulation induces long-term depression (LTD), reducing excitability. Based on this, we hypothesized that 10Hz stimulation would result in a net long-term increase in calcium activity post-stimulation, with the strongest effect observed in cells activated during the stimulation. Conversely, we predicted that 1 Hz stimulation would lead to a net long-term reduction in calcium activity at the population level, with the greatest suppression occurring in cells inhibited during stimulation.

We first analyzed data at the population level for the 4-time-points post-stimulation using a 2-way ANOVA **(Figure 3A)**. We observed a significant effect of stimulation group (F_(3,13724)_=30.97, *p*<*0*.*0001*) but no significant effect of time (F_(3, 13724)_=0.23, *p*=0.87) nor a significant group × time interaction (F_(9, 13724)_=0.2, *p>0*.*9*). Post hoc analysis of the overall group effect showed that 10Hz stimulation was not different than sham (*p>0*.*9*), while 1 Hz stimulation induced a significant inhibition of glutamatergic cells compared to both sham 1 Hz (*p < 0*.*0001*) and 10 Hz stimulation (*p < 0*.*0001*) (**Figure 3B**). Next, we examined the activity of specific subpopulations as noted above. In excited cells, there was a significant effect of stimulation (F_(3,6080)_=24.45, *p<0*.*0001*), but no effect of time (F_(3, 6080)_=0.85, *p*=0.46) or group ×time interaction (F_(9, 6080)_=0.44, *p>0*.*9*). Contrary to our hypothesis, post hoc tests demonstrated that both 10 Hz and 1 Hz stimulation led to significant reductions in activity compared to their respective sham controls (*p<0*.*0001* and *p<0*.*0001* respectively; **Figure 3C, D**). Notably, there was no significant difference in post-stimulation activity between 10 Hz and stim 1 Hz conditions (*p>0*.*9*). In inhibited cells, we again found a significant effect of stimulation (F_(3,7628)_=11.1, *p<0*.*0001*), but no effect of time (F_(3, 7628)_=0.29, *p*=0.83) nor a group × time interaction (F_(9, 7628)_=0.14, *p>0*.*9*). Post hoc analysis revealed that 10 Hz stimulation resulted in reduced suppression (i.e. greater than expected activity) of inhibited cells compared to sham 10 Hz (*p = 0*.*0052*), whereas 1 Hz stimulation produced increased suppression relative to sham 1 Hz (*p = 0*.*04*; **Figure 3E, F**). Furthermore, 1 Hz stimulation showed significantly greater suppression compared to 10 Hz (*p < 0*.*0001*), indicating that the inhibitory effect was more pronounced following 1 Hz stimulation.

**Figure 3.**
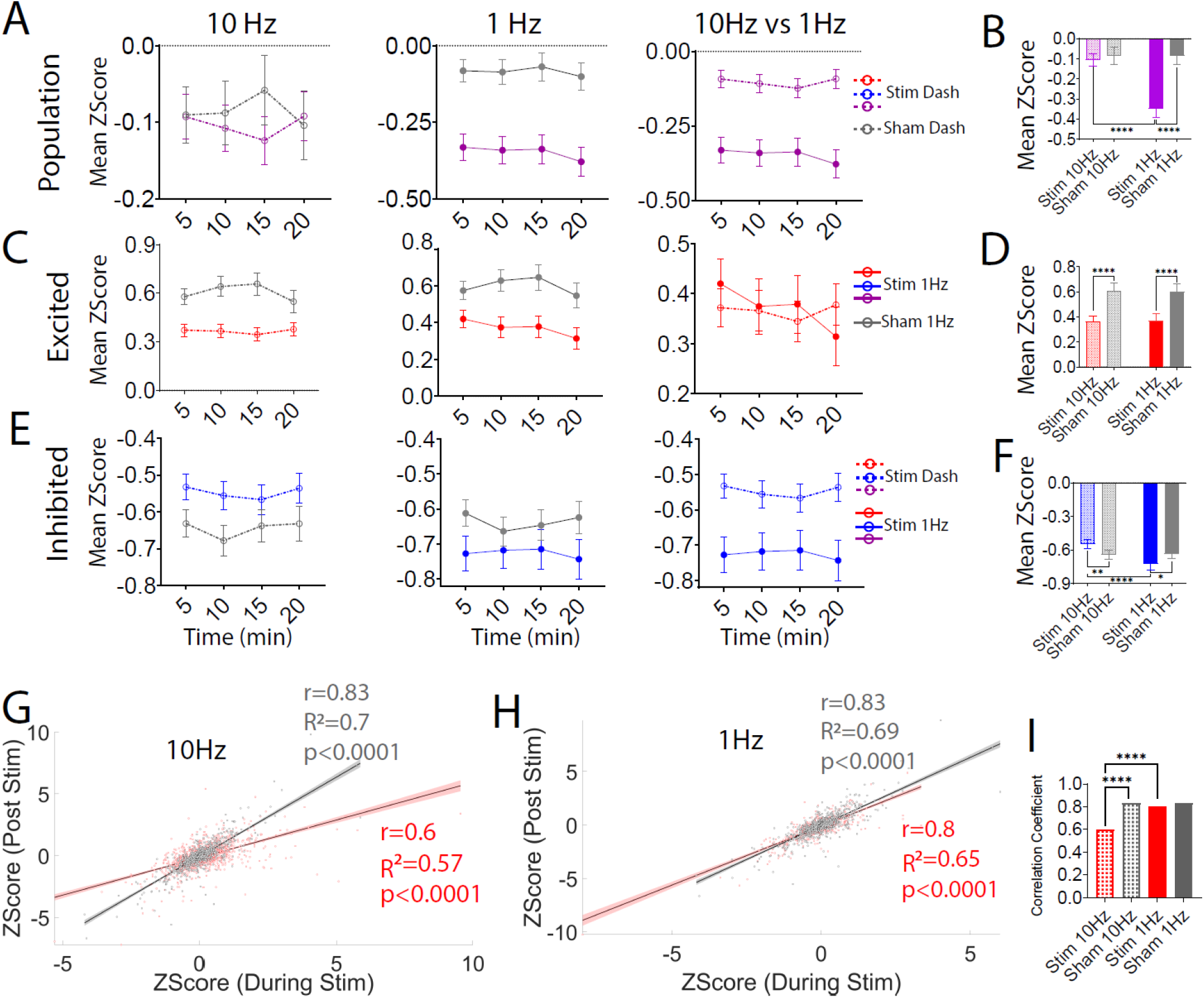
CaMKII Neuronal Responses Post Stimulation. **(A)** Mean Z-score of overall cellular activity during the post-stimulation period across different stimulation protocols and sham conditions for the entire population. Sample sizes: stim 10 Hz (n = 1156), sham 10 Hz (n = 860), stim 1 Hz (n = 562), and sham 1 Hz (n = 857). **(B)** Statistical analysis of mean Z-scores across all post-stimulation time points revealed no significant differences between stim 10 Hz and sham 10 Hz. In contrast, stim 1 Hz resulted in significant inhibition compared to sham 1 Hz (p < 0.0001) and stim 10 Hz. **(C)** Mean Z-score of post-stimulation activity for excited cells. Sample sizes: stim 10 Hz (n = 562), sham 10 Hz (n = 365), stim 1 Hz (n = 194), and sham 1 Hz (n = 383). **(D)** Statistical analysis indicated a significant decrease in excited cell activity for both stim 10 Hz and stim 1 Hz compared to their respective shams (p < 0.0001), but no significant differences between the two stimulation protocols. **(E)** Mean Z-score of post-stimulation activity for inhibited cells. Sample sizes: stim 10 Hz (n = 594), sham 10 Hz (n = 475), stim 1 Hz (n = 368), and sham 1 Hz (n = 474). **(F)** Statistical analysis showed reduced suppression of inhibited cells post stim 10 Hz compared to sham 10 Hz (p = 0.0052) and increased suppression post stim 1 Hz compared to sham 1 Hz (p = 0.04). Stim 1 Hz also showed significantly greater suppression compared to stim 10 Hz (p < 0.0001). All data are presented as mean ± SEM. Statistical analyses were conducted using two-wayANOVA followed by Sidak’s correction for multiple comparisons. Significance levels: *p < 0.05, **p < 0.01, ***p < 0.001, ****p < 0.0001. **(G)** Scatter plot indicating mean Z-score activity during and post stimulation. Linear regression analysis showed a reduced R-squared value for stim 10 Hz, indicating decreased model fitness compared to sham 10 Hz. **(H)** Stim 1 Hz did not result in noticeable changes in R-squared values compared to sham 1 Hz. **(I)** Correlation coefficients were significantly lower for stim 10 Hz compared to sham 10 Hz (p < 0.0001) and stim 1 Hz (p < 0.0001). Data are presented as mean ± SEM. Correlation coefficients and one-way ANOVA were used for statistical comparisons. Significance levels: *p < 0.05, **p < 0.01, ***p < 0.001, ****p < 0.0001.

Thus, results for 1Hz stimulation demonstrated a generalized and consistent reduction in calcium activity across both excited and inhibited cells, explaining the strong inhibition observed at the population level. By contrast, results of 10Hz stimulation demonstrated opposing effects (lower than expected calcium activity in excited cells and greater than expected calcium activity in inhibited cells), an effect that in total cancels each other out, thus demonstrating a null effect at the population level. We also we measured whether there was a difference in the overall distribution of cells that were excited/inhibited post-chi-square analysis on the proportions of excited, inhibited, and non-responsive cells during the 20-minute post-stimulation window. Significant proportional changes were observed for excited cells (χ ^2^ (3) = 12.7, p = 0.005) and inhibited cells (χ ^2^ (3) = 20.31, *p < 0*.*0001*), with 1 Hz stimulation leading to a decrease in the percentage of excited cells and an increase in inhibited cells. No significant changes were noted for non-responsive cells (χ ^2^ (3) = 5.12, *p = 0*.*16*). These findings suggest that 1 Hz stimulation not only induces sustained suppression but also shifts the balance in the overall distribution of cells that are excited vs inhibited post-stimulation. There were no significant differences in the distribution of excited vs inhibited cells post-stimulation for 10Hz compared to sham (See table S2 for all post hoc comparisons).

After observing that 1 Hz stimulation led to prolonged suppression and proportional changes in neuronal activity, while 10 Hz enhanced excitability during stimulation but failed to sustain these effects post-stimulation, we sought to investigate the relationship between changes in activity during vs. post-stimulation. **Figure 3G** illustrates the linear regression analysis comparing neuronal activity during and after stimulation. For 10 Hz stimulation, the analysis revealed a reduced R-squared value compared to sham 10 Hz, indicating a weaker linear relationship between activity during and after stimulation. Furthermore, 10 Hz stimulation resulted in significantly lower correlation values compared to both sham 10 Hz (p < 0.0001) and 1 Hz stimulation (p < 0.0001; **Figure 3I**). By contrast, 1 Hz stimulation did not produce noticeable changes in R-squared values when compared to sham (**Figure 3H-I**). These results are consistent with are prior findings that cells activated during 10Hz stimulation showed lower than expected calcium levels post-stimulation, while the opposite was observed for cells that were inhibited during stimulation.

### Stimulation at 10 Hz Causes Post-Stimulation Suppression of mDLX Neurons

Our investigations of calcium levels following 1Hz and 10Hz neurons was motivated to better understand effects observed clinically in humans that 10Hz stimulation can increase excitability while 1Hz stimulation decreases excitability of frontal cortex in humans. As 10Hz stimulation did not clearly increase calcium activity of glutamatergic neurons at a population level, we next explored whether this stimulation affected GABAergic neurons. Prior studies, in both humans and animals, have suggested that GABAergic cells play a key role in modulating cortical excitability (including stimulation-evoked motor map plasticity^8,11,22,23^). We thus hypothesized that 10 Hz/high-frequency stimulation reduces the intrinsic excitability of GABAergic cells, thereby increased excitability.

To assess this, we performed a similar set of experiments as described above but in animals in which GCAMP8f was expressed in mDLX neurons thereby enriching expression in GABAergic cortical neurons (**Figure 4A)**. We first performed a two-way ANOVA comparing means of calcium in 4 stimulations groups × 6 timepoints (including baseline, stimulation and 20 min post stim in 5 min time bins). We found a significant group effect, (F_(3,6114)_=15.94, *p<0*.*0001*), significant effect of time (F_(5,6114)_=11.91, *p<0*.*0001*) and significant interaction (F_(15,6114)_=2.02, *p=0*.*01*; **Figure 4B**). Post hoc analysis indicated that neither 10 Hz nor 1 Hz stimulation produced significant changes in calcium activity during stimulation compared to sham conditions (p>0.9 for both; **Figure 4B and C**), suggesting that mDLX neurons may be less responsive to stimulation phase compared to CaMKII neurons.

**Figure 4.**
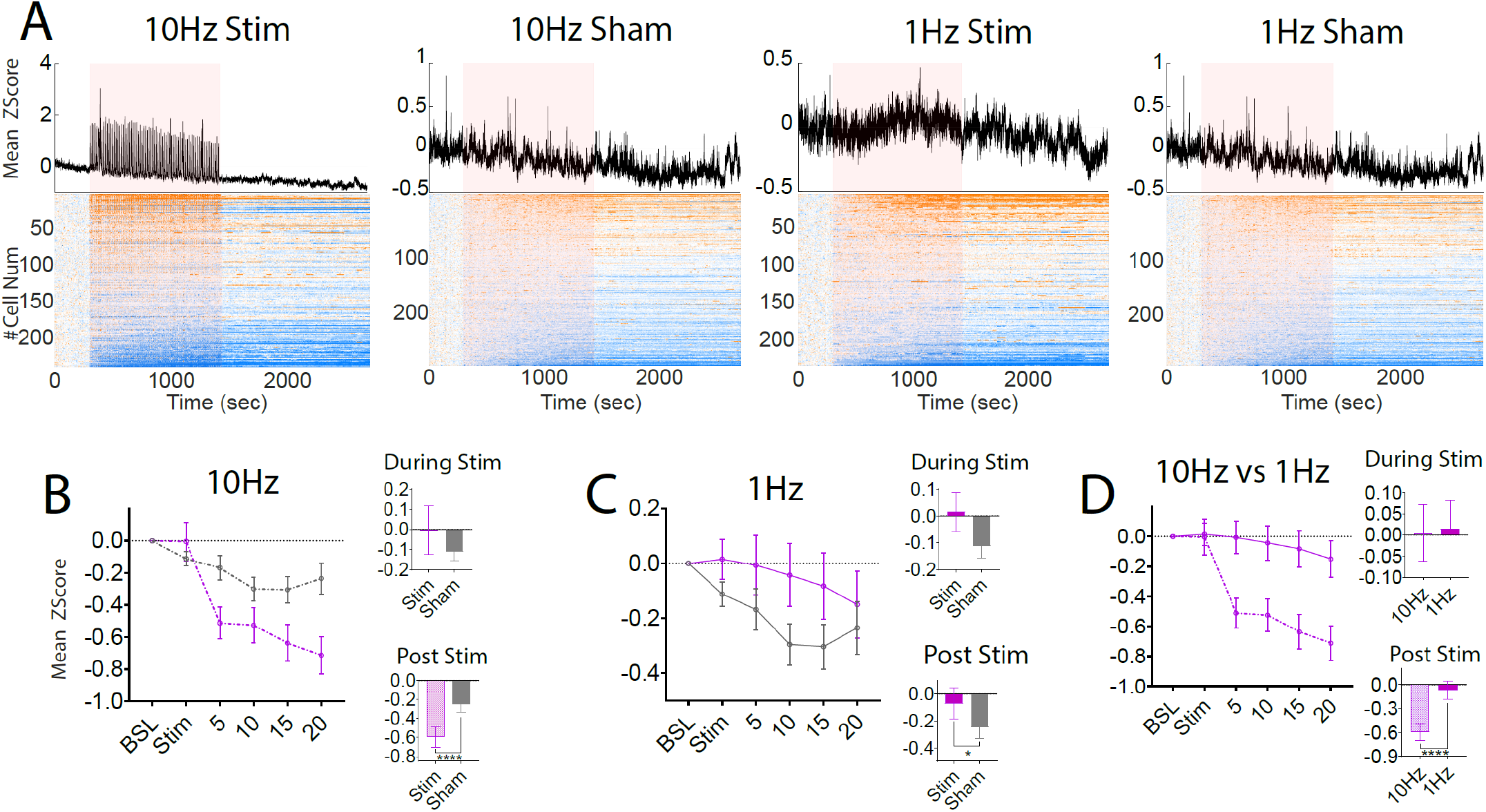
mDLX Neuronal Responses During and After Stimulation. **(A)** Average Z-score of overall cellular activity throughout the entire recording session for different stimulation protocols and sham conditions. Z-scores were calculated based on a 300-second baseline period prior to stimulation. **(B-D)** Mean Z-score of overall cellular activity for different stimulation protocols and sham conditions among the entire population. Sample sizes: stim 10 Hz (n = 219), sham 10 Hz (n = 292), stim 1 Hz (n = 221), and sham 1 Hz (n = 291). During stimulation, no significant changes were observed. Post-stimulation analysis revealed significant suppression in stim 10 Hz compared to sham (p < 0.0001) and stim 1 Hz compared to sham (p < 0.0001). 1 Hz stim showed significant decrease in suppression compared to sham (*p=0*.*04*). between group comparison indicated a significant suppression in stim 10 Hz compared to stim 1Hz. Data are presented as mean ± SEM. Statistical analyses were performed using two-way ANOVA followed by Sidak’s correction for multiple comparisons. Significance levels: *p < 0.05 and ****p < 0.0001.

We next analyzed the post-stimulation activity of mDLX neurons to determine if there were any post-stimulation effects. We found that 10 Hz led to significant suppression of overall neuronal activity compared to sham (p<0.0001) while 1Hz stimulation resulted in significantly less suppression of neuronal activity compared to sham (p=0.04, **Figure 4C**). Moreover, 10 Hz stimulation resulted in a significantly greater suppression in calcium activity of GABAergic neurons compared to 1 Hz stim (*p<0*.*0001*; **Figure 4D**).

## Discussion

In this manuscript we were interested in developing a more mechanistic understanding of how clinical protocols (1Hz and10Hz rTMS) affect cellular activity and plasticity in glutamatergic and GABAergic neurons in prefrontal cortex. Figure 5 summarizes our basic findings: 10Hz stimulation seems to result in a greater long-term suppression of GABAergic activity (relative to glutamatergic cells, in comparison to 1Hz and sham), which could explain changes in stimulation evoked responses observed in humans. By contrast, 1Hz stimulation seems to result in a greater long-term suppression of glutamatergic cell activity (relative to GABAergic cells, in comparison with 10Hz stimulation and sham). This suggests that 10Hz and 1Hz can bidirectionally modulate excitability within prefrontal cortex but via very different mechanisms. Effects on calcium activity during stimulation were only partially predictive of post-stimulation effects (for 1Hz stimulation of glutamatergic cells), suggesting simplistic models of how longer-term changes to excitability occur need to be revisited.

**Figure 5.**
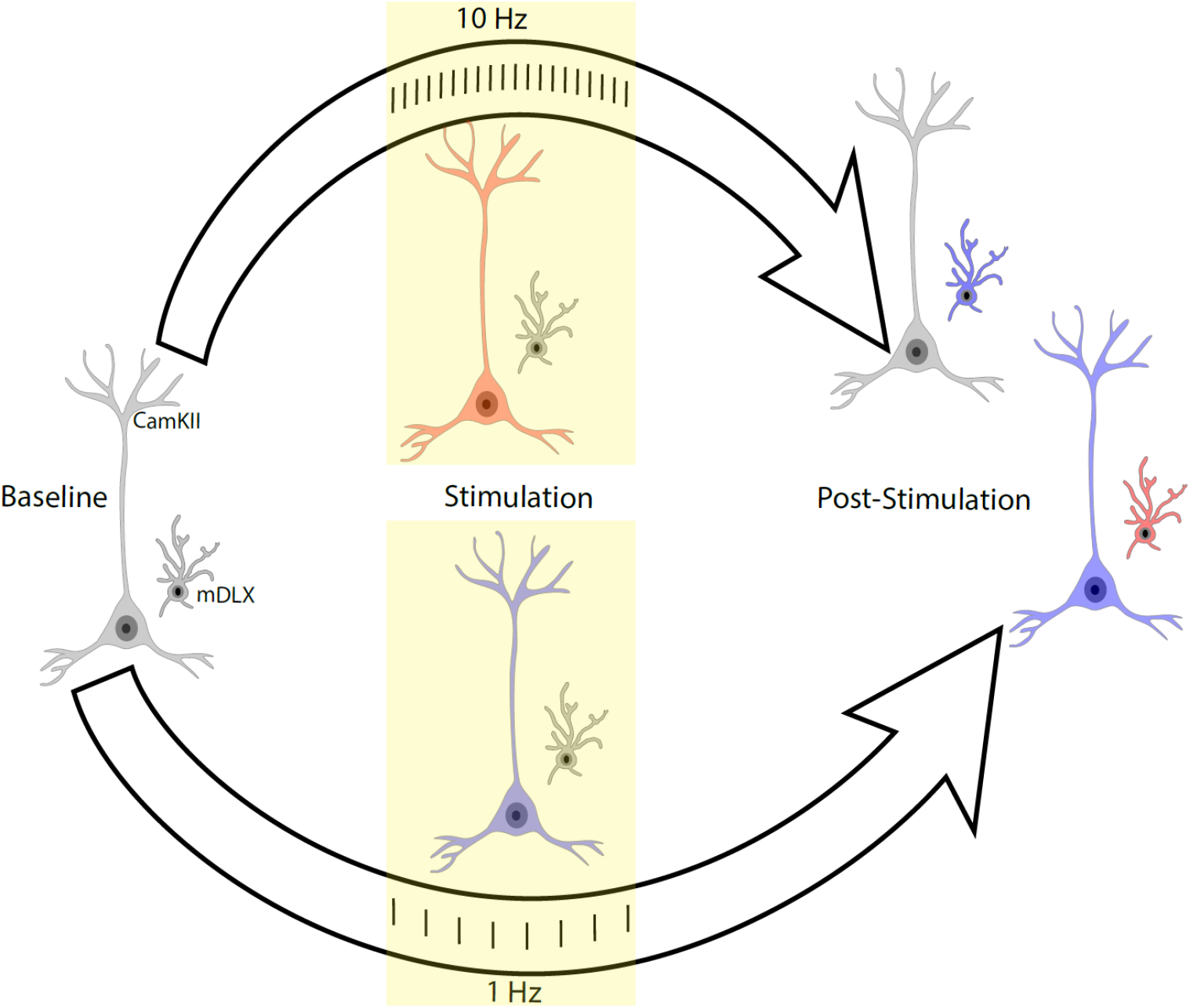
Conceptual Map of Stimulation Effects. The conceptual map visualizes the effects of stimulation protocols on neuronal populations. Colors indicate normalized activity (stim/sham) for the entire population. Gray represents baseline activity, while red and blue depict increased and decreased calcium influx, respectively, relative to sham. Cells with elongated morphology represent CaMKII neurons, and shorter cells represent mDLX neurons. The yellow-shaded area marks the stimulation period. **During Stimulation:** Stim 10 Hz caused increased activity in CaMKII neurons, with no changes observed in mDLX neurons. In contrast, stim 1 Hz induced suppression in CaMKII neurons, while mDLX neurons remained unaffected. **Post-Stimulation:** Stim 10 Hz had no effect on CaMKII neurons but caused suppression in mDLX neurons. On the other hand, stim 1 Hz induced suppression in CaMKII neurons and increased activity in mDLX neurons. These results suggest that different stimulation frequencies selectively modulate neuronal subtypes, with CaMKII neurons showing frequency-dependent excitability changes during stimulation, and mDLX neurons exhibiting delayed, post-stimulation responses.

By recording from individual cells both during and post-stimulation, our work can shed light on one of the fundamental theorized mechanisms by which high-frequency brain stimulation is supposed to work: LTP. According to the LTP model, high-frequency stimulation of prefrontal cortical neurons results in an NMDA-dependent influx of calcium that triggers signaling cascade molecules that increase AMPA expression in synapses ^2,3^. We did, indeed, find that there was an influx of calcium in a certain sub-population of cells during 10Hz stimulation (“excited cells”). Surprisingly, these cells did not show significant long-term changes in excitability post-stimulation, rather they actually showed a reduction in spontaneous calcium activity post-stimulation compared to sham stimulation. Conversely, we found that even though GABAergic neurons were not strongly modulated during 10Hz stimulation, they showed a strong reduction in calcium activity in the post-stimulation period. Given these cells were not directly modulated during stimulation, this suggests that the effects of 10Hz on GABAergic neurons may be indirect and mediated via some network-level plasticity. It is possible that some of the effects we observed may be explain in a cortical network model. For example, if cells excited during the 10Hz protocol that were suppressed post-stimulation preferentially act on GABAergic neurons, this reduced drive could explain reduced activity of these GABAergic neurons. Importantly, there is prior evidence that excitatory rTMS protocols lower GABAergic excitability as assessed using paired-pulse stimulation protocols^11^.

We found that low-frequency stimulation results in a strong reduction in glutamatergic cell activity both during and then after the stimulation ends, and this effect lasts at least 20 minutes and interestingly, in less suppression of GABAergic neurons than would be expected in the sham group. The classic model of 1Hz stimulation is that low-frequency stimulation results in a moderate level of calcium influx which, when uncoupled with a lot of activity in the cell, drives intracellular signaling cascade pathways that result in LTD^3,24^. Our data is far more complicated. In glutamatergic neurons, at a population level, we observed a mean reduction in calcium levels during stimulation and this reduction is sustained. Even when split into both “excited’ and “inhibited” populations, we observed post-stimulation reductions in calcium activity in both sub-populations. Thus, our data does not, clearly show what we would have expected according to the classic/canonical model of LTD, even though we observe a reduction in calcium activity post-stimulation as the model would have predicted. This, again, suggests two possibilities: either there are both direct and indirect/network level effects being observed here; or else calcium sensors are possibly too slow or otherwise not sensitive at detecting some of the stimulation-evoked changes in calcium influx that drive changes observed.

There are several important limitations of this work. First, we are measuring only one aspect of activity (spontaneous calcium levels within the cell), which may or may not reflect other important aspects of plasticity including, for example, inhibitory/excitatory post synaptic potentials; spiking levels; and stimulation-evoked responses. Thus, there may be many other changes that are induced in cells as a result of these protocols that our study was not sensitive to detect. Relevantly most studies in humans use stimulation-evoked responses as a marker of changes in excitability which we did not do. Second, the calcium imaging sensors may affect LTP/LTD processes as they act as a calcium buffer and thus may complicate interpretation. It is possible that cells may act differently if they do not have calcium sensors. Third, we used slightly different calcium sensors in glutamatergic vs. GABAergic neurons. Different sensitivities/kinetics may contribute at least partially to differences observed. This will need to be clarified in future work. Fourth, we use electrical stimulation but are trying to contribute to knowledge regarding magnetic stimulation used clinically. We did roughly match our stimulation to what has been used previously in intracortical microstimulation studies^25,26^ and approximates the electric field using in human TMS. More importantly, as we highlighted in the introduction, repetitive TMS studies were themselves based on studies of plasticity using electrical stimulation in animals. Thus, we have no reason to believe that our results would not generalize at least partially to what is done in humans. Finally, given we are studying rodent prefrontal cortex and some of our results seem to rely on network-level effects, it is possible that our results will not generalize to human cortex. It is increasingly the case that human studies are beginning to illuminate effects of TMS using intracortical invasive recordings^27,28^, and thus it may be possible to validate our results at least partially using such recordings in human. Despite these limitations, we believe our results play an important role in contributing to an understanding of how repetitive stimulation affects single cell physiology within prefrontal cortex.

## Acknowledgments

We would like to thank Miranda Koloski for her contributions to data collection. This material is the result of work supported with resources and the use of facilities at the VA San Diego Medical Center and the Center of Excellence for Stress and Mental Health. The contents of this manuscript do not represent the views of the U.S. Department of Veteran Affairs or the United States Government.

---

**Table S1.**
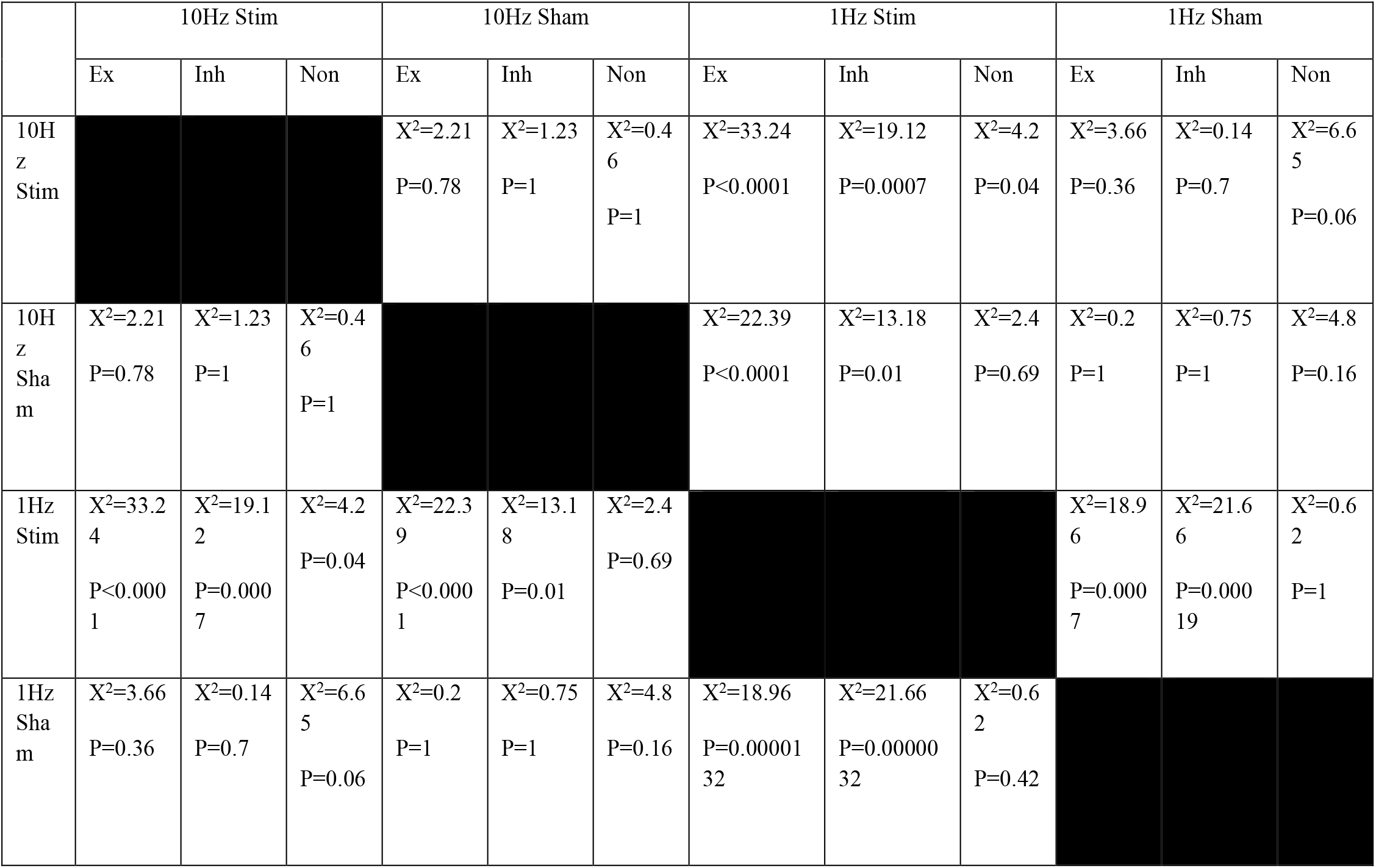
Post hoc multiple comparison for proportion of the cells during stimulation.

|  | 10Hz Stim |  |  | 10Hz Sham |  |  | 1Hz Stim |  |  | 1Hz Sham |  |  |
| --- | --- | --- | --- | --- | --- | --- | --- | --- | --- | --- | --- | --- |
|  | Ex | Inh | Non | Ex | Inh | Non | Ex | Inh | Non | Ex | Inh | Non |
| 10Hz Stim |  |  |  | X<sup>2</sup>=2.21<br>P=0.78 | X<sup>2</sup>=1.23<br>P=1 | X<sup>2</sup>=0.46<br>P=1 | X<sup>2</sup>=33.24<br>P<0.0001 | X<sup>2</sup>=19.12<br>P=0.0007 | X<sup>2</sup>=4.2<br>P=0.04 | X<sup>2</sup>=3.66<br>P=0.36 | X<sup>2</sup>=0.14<br>P=0.7 | X<sup>2</sup>=6.65<br>P=0.06 |
| 10Hz Sham | X<sup>2</sup>=2.21<br>P=0.78 | X<sup>2</sup>=1.23<br>P=1 | X<sup>2</sup>=0.46<br>P=1 |  |  |  | X<sup>2</sup>=22.39<br>P<0.0001 | X<sup>2</sup>=13.18<br>P=0.01 | X<sup>2</sup>=2.4<br>P=0.69 | X<sup>2</sup>=0.2<br>P=1 | X<sup>2</sup>=0.75<br>P=1 | X<sup>2</sup>=4.8<br>P=0.16 |
| 1Hz Stim | X<sup>2</sup>=33.24<br>P<0.0001 | X<sup>2</sup>=19.12<br>P=0.0007 | X<sup>2</sup>=4.2<br>P=0.04 | X<sup>2</sup>=22.39<br>P<0.0001 | X<sup>2</sup>=13.18<br>P=0.01 | X<sup>2</sup>=2.4<br>P=0.69 |  |  |  | X<sup>2</sup>=18.96<br>P=0.0007 | X<sup>2</sup>=21.66<br>P=0.00019 | X<sup>2</sup>=0.62<br>P=1 |
| 1Hz Sham | X<sup>2</sup>=3.66<br>P=0.36 | X<sup>2</sup>=0.14<br>P=0.7 | X<sup>2</sup>=6.65<br>P=0.06 | X<sup>2</sup>=0.2<br>P=1 | X<sup>2</sup>=0.75<br>P=1 | X<sup>2</sup>=4.8<br>P=0.16 | X<sup>2</sup>=18.96<br>P=0.0000132 | X<sup>2</sup>=21.66<br>P=0.0000032 | X<sup>2</sup>=0.62<br>P=0.42 |  |  |  |

**Table S2.**
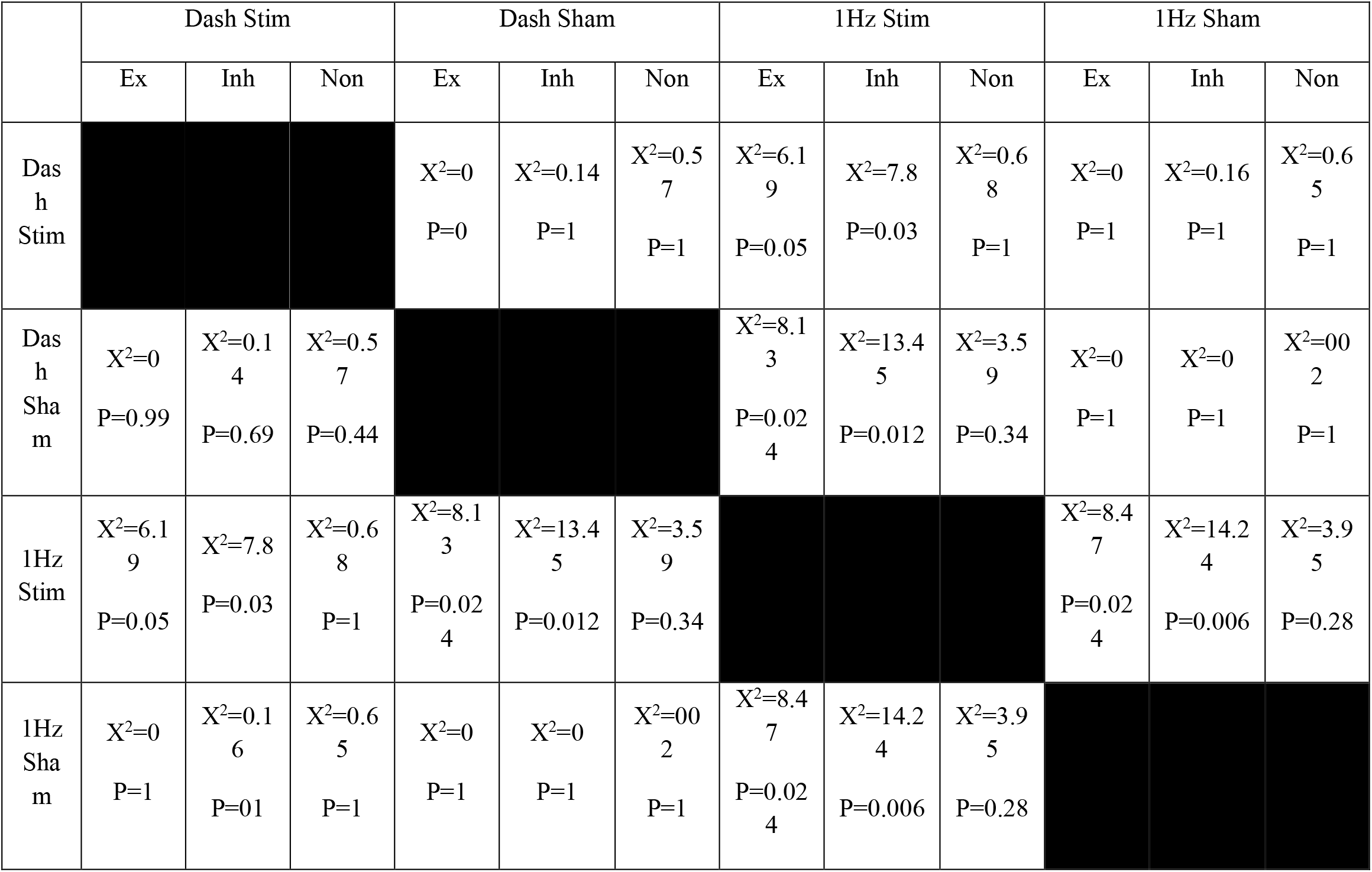
Post hoc multiple comparison for proportion of the cells post stimulation.

